# Transcriptional control of meiotic recombination by the DREAM complex

**DOI:** 10.1101/2025.11.02.686075

**Authors:** Hasibe Tunçay, Xing Ma, Lucas Lang, Franziska Böwer, David Latrasse, Moussa Benhamed, Arp Schnittger

**Affiliations:** Department of Developmental Biology, University of Hamburg, 22609 Hamburg, Germany; Institute of Plant Sciences-Paris-Saclay, Gif-sur-Yvette, France

## Abstract

Meiotic chromosome segregation and recombination, essential for sexual reproduction and evolution, require a specialized gene set. However, how meiotic gene expression is controlled remains largely unclear in multicellular eukaryotes. Here, we identify the DREAM complex as a transcriptional regulator of meiotic recombination in *Arabidopsis*. We show that the DREAM complex localizes to meiotic chromosomes, and that reduced DREAM function disrupts recombination and causes sterility. Meiocyte-specific transcriptomics combined with genome-wide binding analyses revealed that recombination genes, including the crossover-promoting helicase *MER3*, are direct DREAM targets. Notably, bypassing DREAM regulation of *MER3* partially restores recombination in DREAM-deficient plants. Our findings establish the DREAM complex as a major meiotic regulator. Given that reduced DREAM function causes reproductive defects in species such as *Drosophila* and mice, this role may be conserved across multicellular eukaryotes.

## Main

The DREAM complex is an ancient transcriptional regulator across eukaryotes, with conserved components spanning from humans to flowering plants^1,2,3,4,5^. Pioneering studies in *C. elegans, Drosophila* and mammalian cells revealed a conserved and crucial role of the DREAM complex (named after Dimerization partner (DP), RB-like, E2F, and MuvB-core) in regulating genes involved in cell cycle and development^2^. Functionally distinct forms of the DREAM complex have been identified in mammals: the canonical repressive DREAM complex (MuvB, DP, Rb-like, E2F), and the activator complexes MuvB-Myb (MMB) and MuvB-FOXM1^6,7^.

Recently, the full complement of the animal DREAM complex has been identified in the flowering plant *Arabidopsis*, including homologs of the MuvB-core components LIN9 known as ALWAYS EARLY (ALY1, ALY2 and ALY3), the LIN54/TESMIN/TSO1-LIKE CXC homologs (TCX5 and TCX6), homologs of LIN52 (LIN52A and LIN52B), and homologs of ABNORMAL CELL LINEAGE PROTEIN 37 (LIN37A and LIN37B)^3,4,8^. In addition, plant-specific components have been identified^3,9,10^ but it is currently not clear to what extent there are different DREAM/MuvB complexes with specific cell cycle or developmental roles. Functional studies have shown that the *Arabidopsis* DREAM complex transcriptionally represses genes related to the cell cycle and maintenance of DNA methylation^4,8,11^. Additionally, it was found to restrict plant growth following DNA damage^3^.

To explore a possible role of the DREAM complex in reproductive development, we analyzed single mutants of the core MuvB components, i.e., *aly1*, *aly2*, *aly3, tcx5, tcx6, lin37a, lin37b, lin52a*, and *lin52b*. As none of them differed notably from wild-type plants (WT), we generated double mutants of presumed redundant components. While *lin37a/b*, *lin52a/b*, *aly1/3*, and *aly2/3* double mutants were indistinguishable from WT, *aly1/2* and *tcx5/6* mutants exhibited severe sterility, with reduced seed sets (WT, 60 ± 0.4; *aly1/2*, 35±1.8; *tcx5/6*, 11±1.6; Fig. 1A,B) and pollen viability (WT, 99.5%±0.2%; *aly1/2*, 91%±1.0%; *tcx5/6*, 86%±2.2%; *p*<0.0001 for all comparisons against WT; Fig. 1C,D).

**Fig. 1.**
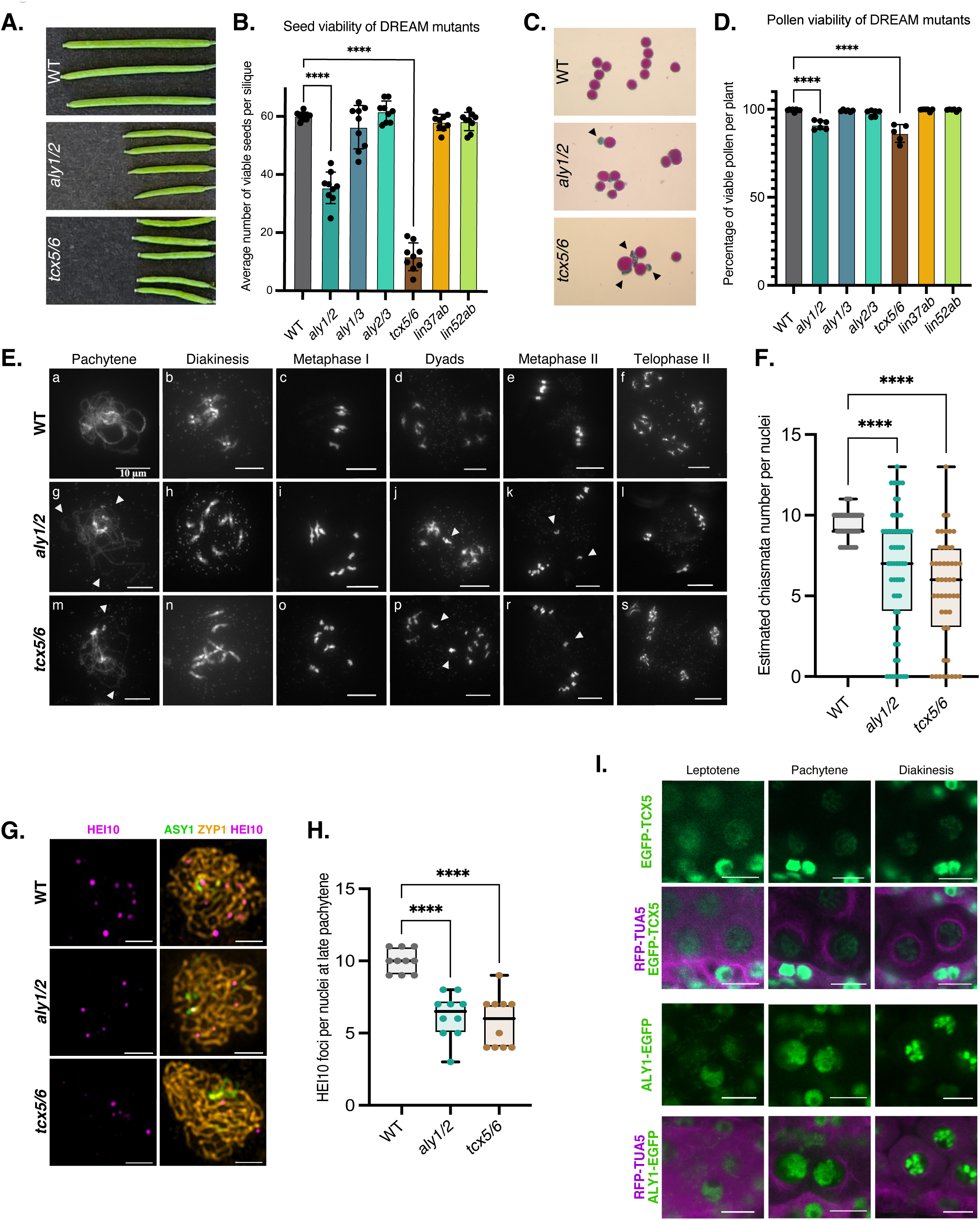
The DREAM complex is vital for successful meiosis. **(A)** The DREAM mutants *aly1/2* and *tcx5/6* show reduced length of siliques. **(B)** Seed viability assessment of mutants of core DREAM components (*n*=10 for all, *p*<0.0001). **(C)** Pollen size variation and death observed in *aly1/2* and *tcx5/6.* Dead pollens are small and slightly blue stained, see black arrowhead. **(D)** Pollen viability assessment for mutants of core DREAM components (*n*=10 for all, *p*<0.0001). **(E)** Meiotic stages in WT (a-f), *aly1/2* (g-l) and *tcx5/6* (m-s) double mutants were observed via chromosome spreads. DREAM mutants have defects in synapsis, crossover formation and chromosome segregation. Representative pictures of pachytene (a,g,m), diakinesis (b,h,n), metaphase I (c,i,o), dyads (d,j,p), metaphase II (e,k,r), and telophase II (f,l,s) are shown for each genotype. White arrowheads indicate pairing defects at pachytene (g,m) and lagging chromosomes at dyad stage (j,p) and metaphase II (k,r). Scale bars, 10 µm. **(F)** Estimated chiasmata numbers were calculated based on metaphase I cells (*n*=40, 47 and 59 for WT, *aly1/2* and *tcx5/6* respectively; *p*<0.0001) **(G)** HEI10 foci at late pachytene in WT, *aly1/2* and *tcx5/6* mutants was observed via immunohistochemical staining of HEI10 (magenta), ASY1 (green) and ZYP1 (orange). Scale bars, 2 µm. **(H)** DREAM mutants exhibit a reduced number of HEI10 foci at late pachytene compared to WT (*n*=10 for all; *p*<0.0001). ONE-WAY ANOVA followed by Dunnett’s multiple comparison test was used for B, D, F, and H. Only significant differences to the WT are indicated by stars. **(I)** ALY1-EGFP and EGFP-TCX5 (both green) expressed in meiocytes and localized to chromosomes; meiocytes in leptotene, pachytene and diakinesis are shown. RFP-TUA5 (magenta) serves as a meiotic stage marker. Scale bars, 10 µm.

Furthermore, both *aly1/2* and *tcx5/6* mutants displayed pollen size variation (Fig. 1C), hinting at gametophytic and/or meiotic defects. However, the transmission efficiency of *tcx6* was unaffected when *tcx5* homozygous/*tcx6* heterozygous plants were crossed with WT (male: 102%, female: 106%, Extended Data Table 1), indicating a sporophytic origin of the defects. Therefore, we analyzed male meiosis by chromosome spreads. Both double mutants showed unpaired chromosomal regions at pachytene (Fig. 1E, g and m, arrowheads), whereas WT chromosomes were fully paired. Pairing defects are followed by incomplete synapsis at late pachytene (Extended Data Fig. 1A). At diakinesis and metaphase I, univalents were observed in the mutants (Fig. 1E, h, n, i, o), instead of the five bivalents seen in WT (Fig. 1E, b and c). The average number of bivalence was 5.0 in WT, 3.8 in *aly1/2* and 3.3 in *tcx5/6* (*n*=81, 48, and 46, respectively; aberrant metaphase I cells showing chromosome entanglements were excluded from this calculation). At interkinesis, we identified lagging chromosomes and unbalanced chromosome pools (Fig. 1E, j, p, k, r, white arrowheads) and after meiosis II the formation of triads and polyads could be observed (Fig. 1E, l and s, Extended Data Fig. 1B), instead of balanced tetrads as in WT (Fig. 1E,f).

These meiotic abnormalities indicated severe defects in recombination in DREAM complex mutants. Indeed, the average number of chiasmata, the physical manifestation of crossovers (CO), during metaphase I (Fig. 1F), was significantly reduced in both *aly1/2* (6.5±0.5) and *tcx5/6* (5.3±0.5) compared to WT (9.5±1.7; *n*=59, 47, 39; *p*<0.0001 for all comparisons against WT). Notably, 11% of *aly1/2* and 15% of *tcx5/6* meiocytes lacked chiasmata entirely, demonstrating that the DREAM complex is important for CO formation and faithful chromosome segregation in *Arabidopsis*.

To determine which recombination step was impaired in DREAM mutants, we assessed meiotic double-strand break (DSB) formation. DSBs are catalyzed by the SPO11 complex^12,13^ in the context of the chromosome axis, a proteinaceous structure, from which DNA loops emerge. The axis includes integral axis proteins such as ASYNAPTIC3 (ASY3) as well as axis associated proteins such as ASY1^14,15^ which build a scaffold for SPO11. SPO11-mediated DSBs generate 3’ single-stranded DNA (ssDNA) overhangs coated by RAD51 and DMC1 to promote strand invasion and joint molecule (JMs) formation^16,17^. Quantification of RAD51 foci at zygotene revealed no significant difference between *aly1/2*, *tcx5/6*, and WT (*p*=0.64 for *aly1/2* vs WT and *p*=0.96 for *tcx5/6* vs WT, One-Way ANOVA; Extended Data Fig. 1C, D), suggesting that DSB formation and their initial processing is unaffected in DREAM complex mutants.

A subset of processed DSBs is resolved as COs by two pathways: class I and class II^18–20^. Class I COs, which account for ∼85% of total COs in *Arabidopsis*, rely on the so-called ZMM proteins such as MER3^21^, PTD^22^, SHOC1^23^, ZIP4^24^, MSH4/5^25^, and HEI10^26^. Assessing class I COs by quantifying HEI10 foci in late pachytene, revealed a significant reduction in *aly1/2* (6.2±0.5) and *tcx5/6* (5.8±0.6) compared to WT (10.0±0.2; *n*=10 for all genotypes; *p*<0.0001 for all comparisons against WT; Fig. 1G,H), indicating that DREAM function is in particular required for Class I CO formation. Consistent with a function in meiosis, ALY1 and TCX5 localized on meiotic chromosomes, as revealed by genomic reporter lines, which nearly completely restored fertility in the respective double mutants (Fig. 1I, Extended Data Figs. 2 and 3).

To pinpoint the genes responsible for the meiotic defects in *aly1/2* and *tcx5/6*, we compared the transcriptomes of male meiocytes from both mutant combinations with WT. Whole-transcriptome sequencing yielded over 71 million reads per sample (three replicates per genotype), with >91% of the reads achieving 99.9% base-calling accuracy (Extended Data Table 2). A total of 17,775 genes were identified to be transcribed in WT, 17,840 in *aly1/2*, and 18,110 in *tcx5/6*. Of these, 17,176 genes were commonly expressed across all genotypes (Fig. 2A). Differential expression analysis revealed 655 upregulated and 256 downregulated genes in *aly1/2*, and 1,358 upregulated and 685 downregulated genes in *tcx5/6* compared to the WT (Fig. 2B, C). The greater number and magnitude of gene expression changes in *tcx5/6* (Fig. 2B, C, and Extended Data Table 3) correlate with its more severe meiotic defects relative to *aly1/2* (Fig. 1F, H). Possibly, ALY3 partially compensates for the loss of ALY1/2, consistent with the embryonic lethality of *aly1/2/3* triple mutantsL.

**Fig. 2.**
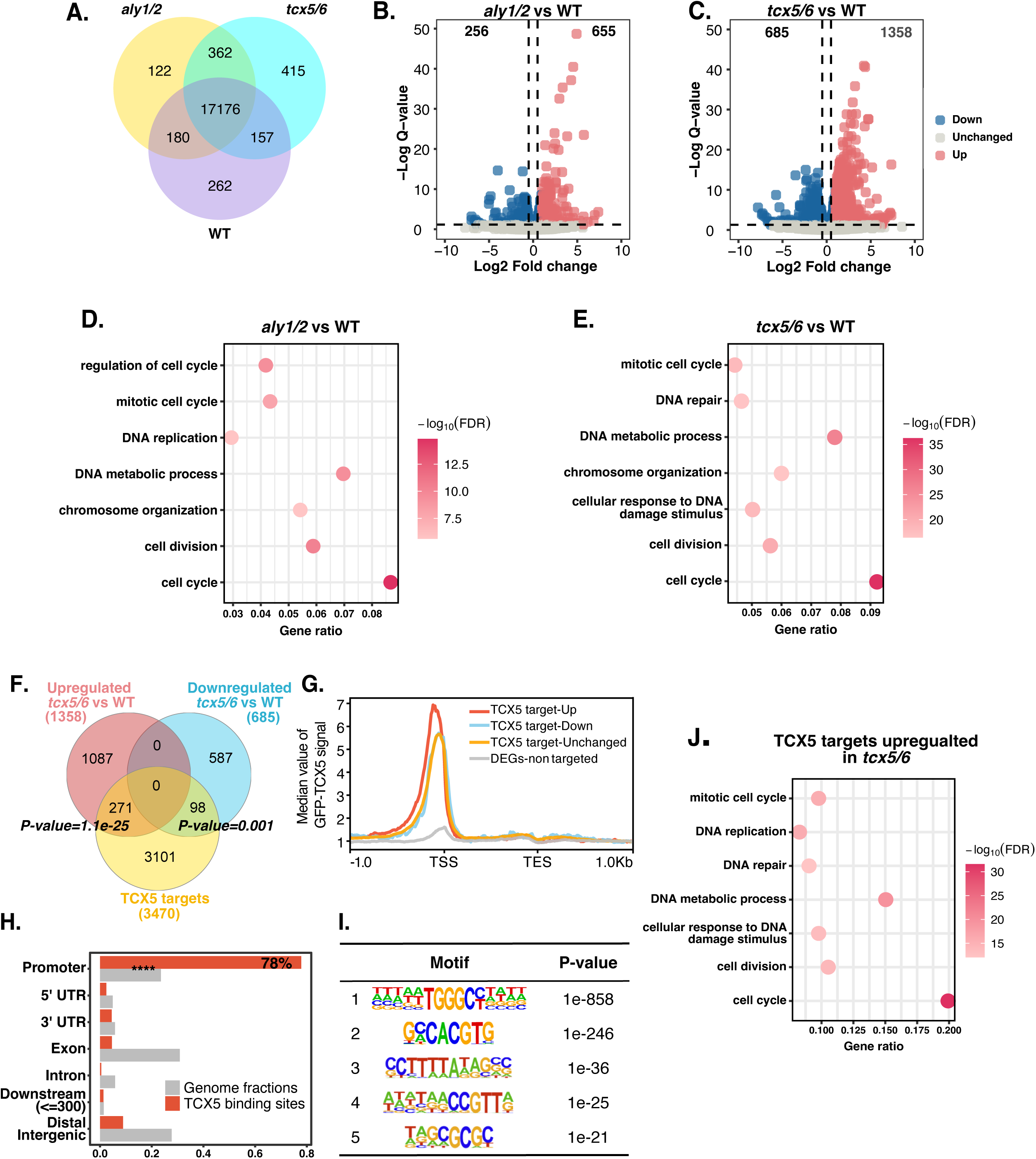
Meiocyte transcriptome analyses of DREAM mutants and TCX5 binding profiles in anthers reveal a role of the DREAM complex in cell cycle regulation and DNA repair in reproductive tissues. **(A)** Venn diagram showing the number of genes expressed in WT, *aly1/2*, and *tcx5/6* male meiocytes. **(B, C)** Volcano plots depicting differentially expressed genes in *aly1/2* **(B)** and *tcx5/6* **(C)** male meiocytes relative to WT. **(D, E)** GO enrichment analysis of upregulated genes in *aly1/2* **(D)** and *tcx5/6* **(E)**, highlighting enrichment in biological pathways related to cell cycle and DNA repair. **(F)** Venn diagram showing significant overlap between upregulated genes (pink), downregulated genes (blue) in *tcx5/6* male meiocytes, and TCX5 targets (yellow) identified by CUT&Tag using EGFP-TCX5 in anthers (Hypergeometric test: *p*=1.1e−25 and *p*=0.0001). **(G)** Meta-plot showing median EGFP-TCX5 CUT&Tag signal across gene bodies. The pink line represents genes upregulated in *tcx5/6*, blue for downregulated genes, yellow for non-differentially expressed genes, and grey for DEGs classified as non-targeted TSS: Transcription Start Site; TES: Transcription End Site. **(H)** Distribution of TCX5 binding sites, showing that 78% of TCX5 targets are bound at promoters (<1 kb from TSS; hypergeometric test, **** *p*<0.0001). Downstream regions are defined as <300 bp from TES; distal intergenic regions are those outside promoter, downstream, and gene body regions; exonic regions include only coding exons. **(I)** HOMER known motif enrichment analysis of TCX5 binding sites at promoter regions showing significant binding motifs and *p*-values. The most enriched motif (1) is GC-rich, followed by a G-box motif (2); motifs recognized by Tesmin/TSO1-like CXC-domain proteins (3), E2Fs (4), and MYB3Rs (5) were also detected. **(J)** GO enrichment of TCX5 targets upregulated in *tcx5/6* male meiocytes.

To identify the biological pathways represented by the differentially expressed genes (DEGs), we performed a gene ontology (GO) enrichment analysis of the deregulated genes in *aly1/2* and *tcx5/6* compared to WT. Among the upregulated genes, both mutants showed a significant enrichment for the GO term *cell cycle*, with 53 genes in *aly1/2* (False discovery rate (FDR)=1.50×10L¹L) and 123 in *tcx5/6* (FDR=5.60×10L³L, Fig. 2D,E), consistent with a conserved role of the DREAM complex in cell cycle regulation^1,7,27^. Additionally, GO terms related to *DNA repair* and *DNA metabolic process* are significantly enriched in *tcx5/6* (62 genes, FDR=3.30×10L¹L and 104 genes, FDR=5.10×10L²L, respectively; Fig. 2D, E). By contrast, GO enrichment analysis of the downregulated genes did not identify any terms directly associated with the meiotic defects observed in the mutants (Extended Data Fig. 4).

To uncover the molecular basis of reduced COs in DREAM mutants, we focused on the expression of genes involved in meiotic recombination. Two observations may account for the CO defects: (i) upregulation of CO-suppressing factors, such as FANCM, RECQ4A, and RMI1, and (ii) downregulation of the CO-promoting factor MER3 (Table 1).

**Table 1.**
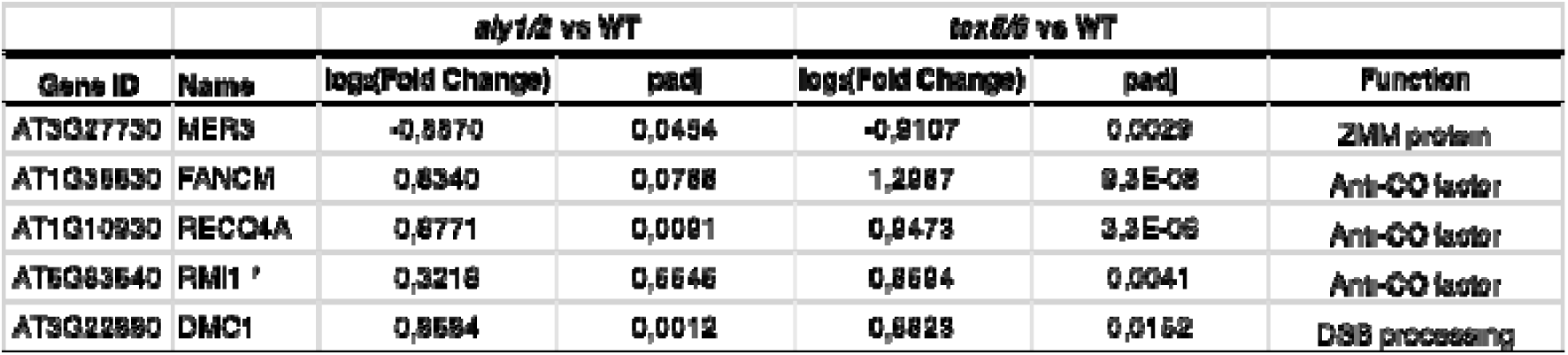
Differential expression of genes in *aly1/2* and *tcx5/6* meiocytes, involved in recombination pathways. For each gene log_2_(Fold-change) values and adjusted *p*-values (*padj*) are provided, together with annotated gene functions. * Deregulation of *RMI* is observed only in *tcx5/6* meiocytes, while its transcript levels are not significantly altered in *aly1/2* compared to WT.

The CO-suppressing factor FANCM promotes DSB repair via synthesis-dependent strand annealing (SDSA)^28^, while RECQ4A/B, being part of the RTR complex together with TOP3 α and RMI1, resolves double Holliday Junctions (dHJ) into non-crossovers (NCOs)^29,30^. Mutants in *FANCM*, *RECQ4* and *RMI1* exhibit elevated CO numbers due to increased class II COs^30,31,32^. Although upregulation of these suppressors has not previously been associated with reduced CO levels, it is tempting to speculate that their elevated expression contributes to the CO defects observed in DREAM mutants.

MER3, a conserved eukaryotic helicase, promotes CO formation by catalyzing D-loop migration and dHJ formation. Like other ZMM mutants, *mer3* exhibits decreased levels of COs leading to univalent formation recapitulating the defects seen in *aly1/2* and *tcx5/6*^33^.

To address whether the transcriptional deregulation of these and other genes in DREAM mutants could be a result of the direct regulation by the DREAM complex, we performed CUT&Tag using anthers of plants harboring a functional EGFP-TCX5 reporter (Extended Data Fig. 3). This resulted in the identification of 3,470 TCX5-enriched genes. Among these, 271 were upregulated and 98 were downregulated in *tcx5/6*, showing a significant overlap between TCX5-enriched targets and differentially expressed genes in the mutant (*p*=1.1e-25 and *p*=0.001, respectively, Fig. 2F).

TCX5 CUT&Tag reads were predominantly enriched around transcription start sites (TSS) as expected for DREAM complex binding^4,34–36^ (Fig. 2G, H; Extended Data Fig. 5A). HOMER motif enrichment analysis of these regions identified several overrepresented motifs, with the most enriched being the *GC rich motif* (*p*=1e-858), followed by a G-Box motif (*p*=1e-246). We also detected motifs recognized by tesmin/TSO1-like CXC domain-containing proteins (*p*=1e-36, Fig. 2I), E2Fs (*p*=1e-21) as well as MYB3Rs (*p*=1e-25; Fig. 2I), which have been shown to be plant DREAM-associated transcription factors^3,4^.

A GO-term enrichment analysis of genes both TCX5-enriched and upregulated in *tcx5/6* revealed significant enrichment for categories related to *DNA repair* and *cell cycle* (Fig. 2J), while TCX5-enriched and downregulated genes in *tcx5/6* did not yield GO terms obviously related to the observed meiotic defects (Extended Data Fig. 5B). Reflecting shared meiotic defects of *aly1/2* and *tcx5/6*, 141 of the 655 genes upregulated in *aly1/2* were direct TCX5 targets, and 119 of these were upregulated in *tcx5/6* (Extended Data Fig. 5C, D). GO term enrichment analysis of these 119 genes similarly highlighted *cell cycle* and *DNA repair* categories (Extended Data Fig. 5E).

Notably, we found no evidence of TCX5 binding at the promoter of *FANCM*, indicating that its upregulation is unlikely to be directly mediated by TCX5. In contrast, TCX5 was detected at the promoter of *RECQ4A* and *RMI1* (Extended Data Table 3). While *RECQ4A* expression was found to be increased in *tcx5/6* and in *aly1/2* meiocytes, *RMI1* transcript levels were only elevated in *tcx5/6* meiocytes (Table 1). In addition, no transcriptional upregulation was observed for *TOP3*α, the third core component of the RTR complex (Extended Data Table 3). These findings suggest that the RTR complex is unlikely to account for the reduced COs observed in plants with impaired DREAM complex activity.

To test whether *MER3* downregulation contributes to reduced recombination observed in DREAM complex mutants, we first confirmed TCX5 enrichment at the *MER3* promoter, suggesting direct regulation (Fig. 3A). To test the relevance of *MER3* downregulation in *aly1/2* and *tcx5/6*, we sought for a way to enhance *MER3* expression in these mutants. We therefore generated a construct in which the genomic sequence of *MER3* fused to *EGFP* was driven by the meiosis-specific *DMC1* promoter. Notably, *DMC1* expression was upregulated in *aly1/2* and *tcx5/6* suggesting that the *DMC1* promoter might confer a strong expression of MER3 in meiocytes (Table 1).

**Fig. 3.**
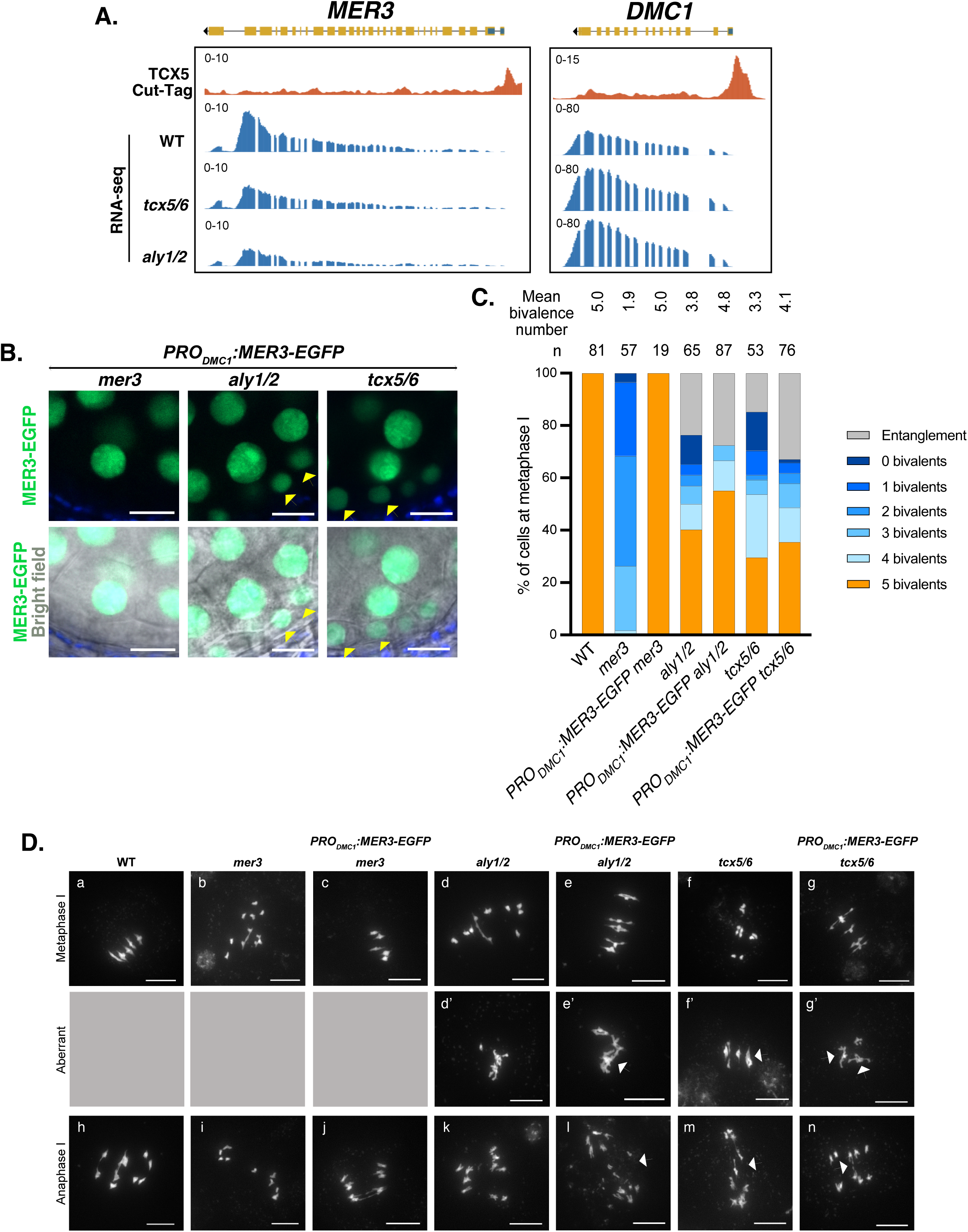
The DREAM complex activates *MER3* transcription relevant for proper CO formation. **(A)** Genome browser snapshot of *MER3* and *DMC1* transcript levels derived from RNA-seq in WT, *tcx5/6*, and *aly1/2* male meiocytes (blue tracks), aligned with CUT&Tag signal for EGFP-TCX5 in anthers (dark orange track). **(B)** MER3-EGFP localization at anthers of *PRO_DMC1_:MER3:EGFP mer3*, *PRO_DMC1_:MER3:EGFP aly1/2*, and *PRO_DMC1_:MER3:EGFP tcx5/6*. The first row shows the MER3-EGFP signal in green and autofluorescence in dark blue. Second row shows a merged image with a bright field to determine cell shape. Yellow arrowheads indicate MER3-EGFP signal at tapetal cells. Scale bars, 10 µm. **(C)** Frequency of observed bivalence numbers and chromosome entanglement at metaphase I. Above the chart, “n” represents the number of metaphase I cells observed and average bivalence number for each transgenic line. Cells showing entanglement were not included in average bivalence number calculation. **(D)** Metaphase I and anaphase I chromosomes displayed for each genotype; WT (a,h), *mer3* (b,i), *PRO_DMC1_:MER3:EGFP mer3* (c,c’j), *aly1/2* (d,d’k), *PRO_DMC1_:MER3:EGFP aly1/2* (e,e’,l), *tcx5/6* (f,f’,m), *PRO_DMC1_:MER3:EGFP tcx5/6* (g,n). White arrowheads mark chromosome entanglement at metaphase I (d’,e’,f’,g’) and chromosome fragmentation at anaphase I (l,m,n). Scale bars, 10 µm.

We observed that MER3-EGFP localization was confined to male meiocytes (Fig. 3B) and, importantly, the *pDMC1-MER3-EGFP* construct fully restored recombination in homozygous *mer3* mutants (Fig. 3C, and Extended Data Fig. 6A), validating its functionality. We next introduced this construct into *tcx5^−/−^ tcx6^+/−^* and *aly1^−/−^ aly2^+/−^* mutants to assess whether increased *MER3* expression could restore CO formation defects. Interestingly, MER3-EGFP was detected not only strongly in meiocytes in *aly1/2* and *tcx5/6* anthers, but also in the surrounding tapetal cells (Fig. 3B, Extended Data Fig. 6B). Given that *DMC1* is a direct TCX5 target (Fig. 3A), one function of the DREAM complex may be to restrict the appropriate spatial activity of the *DMC1* promoter and potentially of other genes involved in meiotic processes to meiocytes.

Strikingly, *MER3-EGFP* expression in *aly1/2* increased the proportion of meiocytes with five bivalents from 40% to 55% (Fig. 3C), and cells exhibiting two, one, or no bivalents were no longer observed. *MER3-EGFP* expression in *tcx5/6* reduced the fraction of cells with no bivalents from 14% to 1% and those with one bivalent from 9% to 4% (Fig. 3C). Furthermore, *MER3-EGFP* expression significantly increased the average bivalent numbers from 3.8 to 4.8 in *aly1/2* (*p*<0.0001) and from 3.3 to 4.1 in *tcx5/6* (*p*<0.008, Fig. 3C and Fig. 3D e, g). However, it also elevated the incidence of chromosome entanglements, potentially contributing to the chromosome fragmentation seen in anaphase I (Fig. 3D l, n). These findings identify *MER3* as a major target of an activating DREAM complex in meiosis.

In animals, the DREAM core complex has been shown to act as both a transcriptional repressor and activator, depending on the developmental context ^34,36^. Our data now reveal a positive, activating function of the DREAM complex in plants, alongside its previously found repressive function. To explore this dual role, we integrated our meiocyte transcriptome data with previously published *tcx5/6* and WT seedling transcriptome data^4^, together with TCX5 binding profiles obtained from seedlings (ChIP-seq)^4^ and anthers (CUT&Tag, this study). We identified 1,011 shared TCX5 target genes, with 2,459 and 1,732 genes uniquely bound in anthers and seedlings, respectively (Extended Data Fig. 7A). Among genes upregulated in *tcx5/6* meiocytes and seedlings, 126 were common TCX5 targets enriched for cell cycle-related GO terms, consistent with the conserved repressive role of the DREAM complex^1,35,37^ (Extended Data Fig. 7B,C). In contrast, TCX5-bound genes showed no significant overlap with genes downregulated in *tcx5/6* seedlings (Extended Data Fig. 7D), whereas a large overlap exists in *tcx5/6* male meiocyte downregulated genes and TCX5-bound genes in anthers (Fig. 2F). Moreover, no downregulated genes in *tcx5/6* meiocytes and seedlings were shared TCX5 targets (Extended Data Fig. 7E). These findings led to the interesting hypothesis that TCX5’s activating role is specific to reproductive tissues.

To further explore the role of the DREAM complex in reproductive development, particularly in meiosis, we examined 135 genes with established meiotic functions that are transcriptionally active in meiocytes (Extended Data Table 3). To this end, we assessed their transcriptional regulation and TCX5 binding in seedlings^4^ and meiocytes/anthers. In seedlings, TCX5 binds to 93 of these genes. Of these, 84 were transcriptionally repressed in DREAM complex mutants, including key regulators of class I and II CO formation, DSB formation and repair, and synaptonemal complex assembly (Extended Data Table 3). These results suggest that the DREAM complex broadly represses meiotic gene expression in non-reproductive tissues, confining their activation to the appropriate developmental context.

To then analyze the transcriptional regulation of these 135 genes in reproductive tissues, we adopted a more permissive cutoff for TCX5 enrichment, i.e., fold-change > 2 versus > 4 used in the genome-wide CUT&Tag analysis. This approach accounts for the low proportion of meiocytes in anther tissue, which may dilute meiocyte-specific signals. Nonetheless, we cannot fully exclude that some TCX5 binding sites unique to meiocytes may remain undetected.

In reproductive tissues, we identified five distinct regulatory patterns that illustrate how the DREAM complex controls reproductive gene expression (Fig. 4A, B, Extended Data Table 3): (I) 14 genes were not bound by TCX5 in neither anthers nor seedlings, indicating that they are not DREAM targets in either context. (II) 16 genes were not bound by TCX5 in anthers but were bound (and either repressed or unregulated) in seedlings, including the ZMM genes *HEI10* and *MSH5*, suggesting their release from DREAM-mediated repression in reproductive tissues. This pattern supports the above observation that the DREAM complex is a key regulator of reproduction-specific gene expression. (III) 59 genes bound by TCX5 in anthers but not differentially expressed in meiocytes, such as *ASY1* and *ASY3* (chromosome axis formation) as well as *SPO11-1* and *SPO11-2* (DSB initiation). Possibly, this indicates a role for the DREAM complex in restricting the expression of these genes to meiocytes within an anther. Thus, loss of DREAM function would not alter their expression in meiocytes, as repression would already be lifted. (IV) 44 genes bound by TCX5 in anthers and repressed in meiocytes, including *RAD51, X-RAY REPAIR CROSS COMPLEMENTING 3 (XRCC3), MEIOSIS DEFECTIVE 1 (MEI1), RMI1,* and *RECQ4A* (Extended Data Table 3). These are largely associated with DNA repair^38,39^, and may be subjected to phase-specific regulation, e.g., during DSB formation and initial repair in leptotene and zygotene. Notably, *DMC1* also falls into this category. As shown above, the loss of DREAM function causes both the ectopic expression of *DMC1* in the anther tissues surrounding the meiocytes and its upregulation within the meiocytes (Fig. 3A, D). Class IV therefore may encompass two regulatory modes: temporal repression (e.g., *DMC1* downregulation after early prophase) and spatial restriction to meiocytes, reflecting the multiple mechanisms through which the DREAM complex governs development-specific gene expression. (V) Two genes, *MER3* and *PS1*, the latter important for proper meiotic spindle formation^40^, were bound by TCX5 in anthers and transcriptionally activated in meiocytes. Given MER3’s essential role in class I CO formation, this highlights an activator function of the DREAM complex in reproductive tissue and suggests a direct link to the control of key meiotic processes.

**Fig. 4.**
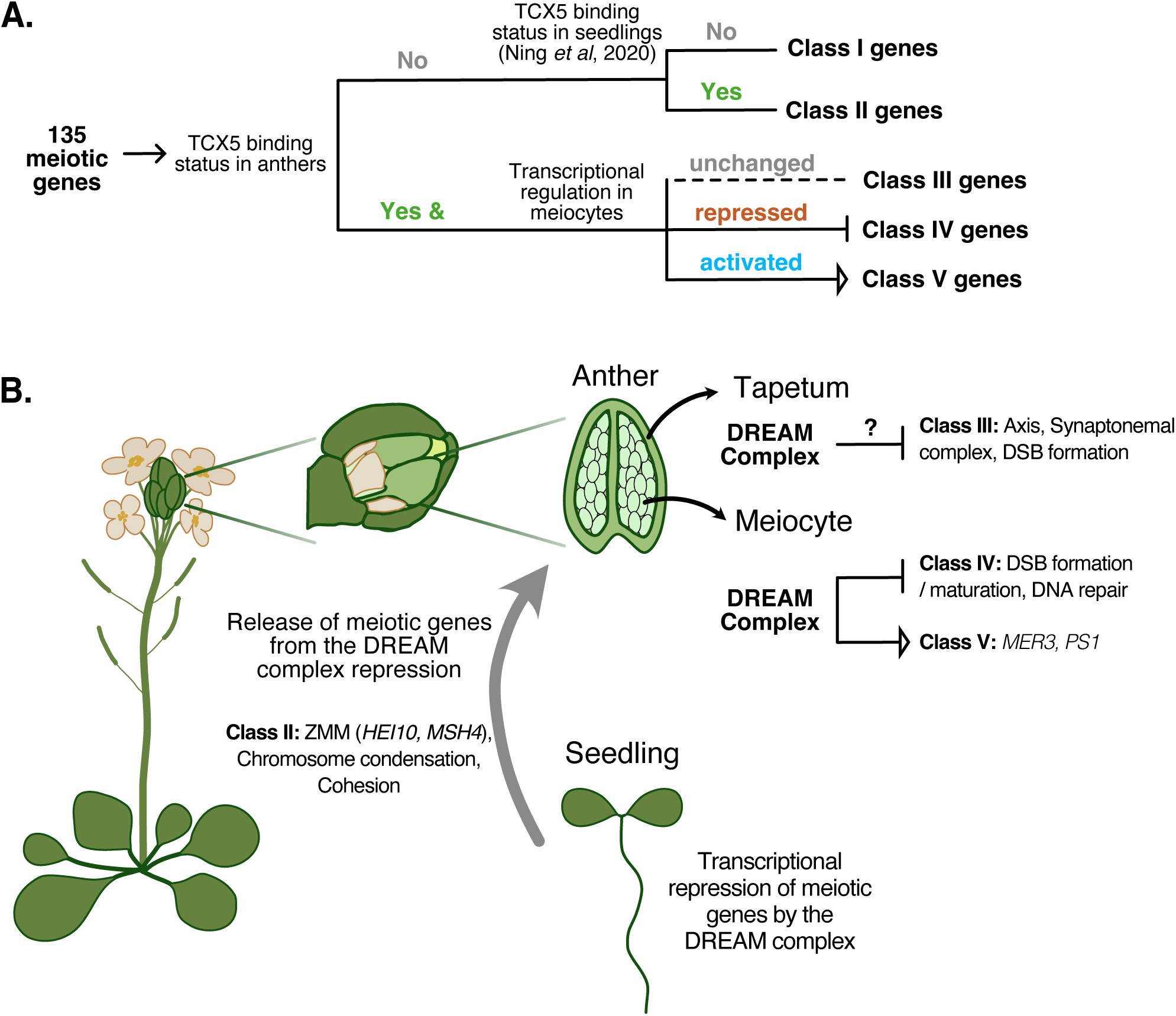
Summary of DREAM complex-mediated regulation of meiotic genes. **(A)** Classification of 135 meiotic genes with established meiotic roles that are expressed in meiocytes, based on TCX5 binding in seedlings and anthers and transcriptional changes in meiocytes. **Class I:** no TCX5 enrichment in either seedlings or anthers, indicating they are not direct DREAM targets. **Class II:** not bound by TCX5 in anthers but bound in seedlings; their transcription is repressed or unchanged in both seedlings and meiocytes. **Class III:** enriched for TCX5 binding in anthers but with no transcriptional changes in meiocytes. **Class IV:** TCX5-bound in anthers and transcriptionally repressed in meiocytes. **Class V:** TCX5-bound in anthers and transcriptionally activated in meiocytes. **(B)** Hypothetical model of DREAM-mediated regulation of meiotic genes across somatic and reproductive tissues. In seedlings, meiotic gene expression is repressed by the DREAM complex. In reproductive tissues, Class II genes are derepressed, indicating reprogramming toward a reproductive state. Class III genes appear regulated at the anther level rather than in meiocytes, potentially restricting their expression to meiocytes. The repression of Class IV genes in meiocytes suggests phase-specific control that confines their expression to a particular stage of meiosis and/or spatial restriction to meiocytes. Class V genes, *MER3* and *PS1*, highlight an unusual activator function of the DREAM complex in reproductive tissue.

Here, we identify the DREAM complex as a major regulator of meiotic gene expression in *Arabidopsis*. A key target of its action is *MER3*. Partial restoration of bivalent formation by MER3 expression, particularly in *tcx5/6*, suggest DREAM governs additional aspects of meiotic recombination. Moreover, the chromosome entanglements seen in particular in *tcx5/6* further implicate DREAM in broader meiotic functions.

Notably, no transcription factor has yet been discovered that directly controls meiotic gene expression in *Arabidopsis* or any other plant species. Our work shows that the DREAM complex functions as both a repressor and an activator of transcription during reproduction. Its repressive role is consistent with prior studies in somatic tissues of *Arabidopsis*^3,4^ and with the conserved function of DREAM complexes in other species^35,37,41^. We demonstrated that the DREAM complex represses a large subset of meiotic genes in somatic cells (e.g., seedlings) mirroring the somatic repression of germline genes observed for dREAM in *C.elegans* ^37,42^. Furthermore, in the reproductive context, the DREAM complex in *Arabidopsis* fine-tunes gene expression within anthers, restricting the expression of recombination genes, including *DMC1,* specifically to meiocytes. Possibly, this repression is even crucial to establish temporal regulation during meiosis, for example, restricting the expression of recombination genes to meiotic prophase I. Importantly, we uncover that DREAM activates key meiotic genes, including *MER3*, and this activator role appears to be specific for reproductive processes raising the possibility of a specialized DREAM variant in reproduction.

Notably, the DREAM complex is required for fertility in different animals, e.g., *C. elegans*, *Drosophila*, and mouse^34,36,43–45^. However, which DREAM regulated genes are responsible for the observed defects have remained unresolved up to now. Thus, our study highlighting the DREAM complex as a critical transcriptional regulator of meiotic genes in plants offers a blueprint for related studies in other organisms.

## Online Methods

### Plant materials

The *Arabidopsis thaliana* accession Columbia (Col-0) was used as a WT reference throughout this study. All mutants and transgenic lines used in this study were in Col-0 background. The T-DNA insertion lines *tcx5-1* (SALK_047165), *tcx6-1* (GK-453H07), *aly1-1* (SALK_073108), *aly2-2* (SALK_118765C), *aly3-4* (SALK_125138C), *lin37a-2* (SALK_057175), *lin37b-3* (SALK_103139C), *lin52b-1* (GK_854A11), and *mer3-2* (SALK_091560) were isolated from GABI-KAT and SALK T-DNA collections and described previously^3^. The *lin52a-c1* mutant is generated via CRISPR/Cas9 and described in our previous study^3^. Transgenic lines harboring *PRO_RPS5A_:TagRFP:TUB5*^46^ and *PRO_ASY3_:ASY3:RFP*^47^ reporter constructs were crossed with EGFP-TCX5 and EGFP-ALY1 transgenic reporter lines generated in this study. All plants were grown in growth chambers with a 16 h light/21°C and 8 h/18°C dark cycle.

### Plasmid construction and plant transformation

To generate *PRO_TCX5_:EGFP:TCX5* reporter construct, genomic sequence of *TCX5* (5557 bp) was amplified by PCR and subsequently cloned into the *pENTR* vector via SLiCE reaction. A *Sma*I restriction site was introduced right after the start codon by PCR. The resulting construct was linearized by *Sma*I restriction and ligated with *EGFP*. To generate the *PRO_ALY1_:ALY1-EGFP* construct, the genomic sequence of *ALY1* (9,252 bp) sequence was amplified as two fragments by PCR and inserted into the *pENTR* vector via SLiCE reaction. A *Sma*I restriction site was introduced right before the stop codon by PCR. The resulting construct was linearized by *Sma*I restriction and ligated with *EGFP*. These constructs were subsequently transferred into the binary expression vector pGWB501^48^ by Gateway LR reaction.

To generate *PRO_DMC1_:MER3:EGFP:TER_REC8,_* the genomic sequence of *MER3* covering the coding sequence (6259 bp) and an *EGFP* fragment were amplified by PCR and incorporated by SLiCE reaction into the *pENTR-PRO_DMC1_:Sma*I*:TER_REC8_*vector linearized by *Sma*I restriction. The resulting fusion construct was then transferred into the binary expression vector pGWB501^48^ by Gateway LR reaction. *Arabidopsis thaliana* transgenic lines were generated by *Agrobacterium-*mediated transformation^49^.

### Pollen and seed viability assessment

Pollen viability was assessed as previously described^50^. 3 open flowers per plant were dipped into 12 µL of Peterson staining solution (10% ethanol, 0.001% malachite green, 25% glycerol, 0.005% acid fuchsin, 0.0005% orange G, 4% acetic acid). Slides were covered with a cover slip and incubated on a pre-heated hotplate at 80°C for 5 min and screened using a light microscope (Axioskop, Carl Zeiss GmbH, Oberkochen, Germany) and the 20× objective. 600 pollen grains of 5 different plants were counted per line. To assess seed viability, seeds were evaluated from 5 siliques of at least 5 different plants per line.

### Cell spreads

Cell spreads of transgenic lines were prepared as previously described^47^. Briefly, fresh flower buds were fixed in ethanol/acetic acid (3:1) for 48 h, at 4 °C and washed with 75% ethanol three times and stored in 70% ethanol at 4 °C. For spreading, flower buds between 0.3 and 0.5 mm in size were digested in an enzyme solution (10 mM citrate buffer containing 1.5% cellulose, 1.5% pectolyase, and 1.5% cytohelicase) for 3 h at 37 °C. Three digested flower buds were transferred onto a glass slide and mashed with a bent needle. Cell suspension was spread on an adhesive slide on a 46 °C hot plate after adding 10 µL of 45% acetic acid. The slide was then rinsed with ice-cold ethanol/acetic acid (3:1) solution and mounted in 15 µL of VECTASHIELD medium with DAPI (Vector Laboratories). Slides were observed using an Axiophot microscope (Zeiss) with 63× oil-immersion objective.

### Immunostaining

Immunostaining was performed as previously described^51^. Briefly, flower buds were dissected using needles, and anthers were transferred to a droplet of artificial pond water (0.5 mM NaCl, 0.4 mM CaCl_2_, 0.2 mM NaHCO_3_, 0.05 mM KCl, in deionized water) on a polylysine-coated slide. Anthers were gently squeezed to release meiocytes. The isolated meiocytes were collected into a 1.5 mL tube on ice and subsequently digested in 15 µL of digestion solution (1% cytohelicase, 1.5% sucrose, and 1% polyvinylpyrrolidone) in the dark for 30 min. Following digestion, meiocytes were transferred to a clean, charged microscope slide. Next, 20 µL of 1% Lipsol was added, and the sample was spread by tilting the slide. After a 4-min incubation at room temperature, 24 µL of 4% formaldehyde was added, and the slide was left to air dry. The slides were washed twice with 2× SCC (0.3 M NaCl and 30 mM Na_3_C_6_H_5_O_7_) for 5 min each. Primary antibody mix was then added, and the slide was covered by a piece of parafilm. Slides were incubated overnight at 4 °C in a humid chamber. Slides were washed twice with 2× SCC for 5 min, followed by addition of the secondary antibody mix. Slides were again covered with parafilm and incubated for 1 h at 37 °C in a humid chamber. Finally, slides were washed twice with 2× SCC for 5 min in the dark. 15 µL of VECTASHIELD medium with DAPI was added to the slide and covered with a glass cover slip and sealed using nail polish. The following dilutions were used for the primary and secondary antibodies: anti-ASY1 1:10,000^51^, anti-RAD51 1:300^52^, anti-ZYP1 1:500^25^, anti-HEI10 1:300^51^, anti-guinea pig Alexa 488 1:400 (Abcam #ab150185), anti-rabbit Alexa 568 1:400 (Abcam #ab175471), anti-rat Alexa 647 (Abcam #ab150159).

### Microscopy

To detect protein localization in reporter lines, young anthers were dissected from floral buds in a drop of artificial pond water (APW) and imaged directly using an upright Zeiss LSM900 confocal microscope equipped with a 60× water-immersion objective. Meiotic stages were determined based on chromosome morphology, cell shape, or microtubule pattern visualized by the *PRO_RPS5A_:TagRFP:TUA5* marker. GFP was excited at 488 nm and detected at 495–560 nm and TagRFP was excited at 561 nm and detected at 570–650 nm.

For imaging immunohistochemical stained samples, high-resolution imaging was performed using a Zeiss LSM 880 confocal microscope equipped with an Airyscan detector (Zeiss, Oberkochen, Germany), using a 63×/1.4 NA oil-immersion objective. Airyscan mode was used to improve resolution and signal-to-noise ratio. Z-stacks were acquired at optimal intervals for 3D reconstruction and processed using the Zeiss Zen Black software (ZEN 2.3 SP1).

### Male meiocyte isolation and RNA extraction

Anthers at developmental stage 6–11 covering all meiotic stages, were dissected and collected in artificial pond water supplemented with 1 U/µl RNasin Ribonuclease Inhibitor (Promega, N2511). Anthers were gently squashed to release meiotic columns, and ∼300 columns per sample were transferred into ZR BashingBead Lysis Tubes (Zymo Research, S6003-50) containing lysis buffer from the RNAqueous™-Micro Total RNA Isolation Kit (Thermo Fisher, AM1931), then immediately flash-frozen in liquid nitrogen. Dissections and collections were performed at 4 °C from 16 plants per genotype for each replicate, across three biological replicates. Samples were homogenized using a mechanical shaker, and total RNA ‘was extracted according to the manufacturer’s protocol. RNA quality was assessed using an Agilent 2100 Bioanalyzer with the NanoChip Kit, and all samples showed high integrity (RIN > 8.0).

### Transcriptome sequencing and data analysis

RNA sequencing libraries were prepared by Novogene and sequenced on an Illumina HiSeq X Ten platform using 150 bp paired-end reads (PE150). Raw read quality was assessed using *FastQC*. The reference genome index was built with *HISAT2* (v2.0.5) and paired-end reads were aligned to the genome using the same version. Gene-level read counts were obtained using *featureCounts* (v1.5.0-p3). Transcript assembly and quantification were performed with *StringTie* (v1.3.3b), and gene expression levels were reported as fragments per kilobase of exon per million fragments mapped (FPKM). Next, we used DESeq2 to identify differentially expressed genes (DEGs) with the thresholds: adjusted *p*-value<0.05 and |log2FoldChange|>0.5. Gene Ontology (GO) enrichment analysis of DEGs was performed using agriGO ^53^ v2.0 (https://systemsbiology.cau.edu.cn/agriGOv2/). GO terms with an adjusted *p*<0.05 were considered significantly enriched.

### CUT&Tag sample preparation from Arabidopsis anthers

Anthers at stage 6–11 from the *PRO_TCX5_:EGFP:TCX5 tcx5* transgenic line were dissected to capture all stages of meiosis and collected in Nuclei Extraction Buffer (0.25 M sucrose, 10 mM Tris-HCl pHL8.0, 10LmM MgClL, 1% Triton X-100) supplemented with a protease inhibitor cocktail (cOmplete™ ULTRA Tablets, Roche, 05892791001). Anthers were homogenized using a glass Dounce tissue grinder and incubated on a rotating wheel at 4 °C for 10 min. The homogenate was filtered through 50 µm filters and nuclei were pelleted by centrifugation at 1,000L×Lg for 10Lmin at 4 °C. Nuclei were washed three times with the same extraction buffer. The final pellet was resuspended in sucrose buffer (0.25 M sucrose, 10LmM Tris-HCl pHL8.0, 10LmM MgClL) containing a protease inhibitor cocktail (cOmplete™ ULTRA Tablets, Roche, 05892791001) and 10% DMSO. Extracted nuclei were cryopreserved at −80 °C in a controlled-rate freezing container. CUT&Tag was performed using the CUT&Tag-IT® Assay Kit, Anti-Rabbit (Active Motif, 53160), with an anti-GFP antibody (ThermoFisher, A-11122) used to target EGFP-tagged TCX5. Library preparation was carried out according to the manufacturer’s protocol.

### CUT&TAG data analysis

Paired-end reads of CUT&Tag raw data were trimmed by using Fastp (v0.23.2). Using Bowtie2^54^ (v2.5.4) the cleaned reads were aligned to the Arabidopsis reference genome assembly TAIR10. Duplicated reads were removed and reads were filtered to keep only high quality alignments (MAPQ score>30) using SAMtools^55^ (v1.9) Afterwards, peaks were called using MACS3^56^ (https://github.com/macs3-project/MACS) with the threshold *q*-value<0.05. ChIPseeker from the R package was used to investigate genomic distribution of CUT&Tag peaks. For genome-wide analyses, TCX5 targets were defined as regions with a fold-enrichment > 4. For analyses focusing on the promoters of the 136 meiotic genes (Extended Data Table 3), the fold-enrichment threshold was set to > 2 to reduce the possibility of false negatives, given the low proportion of meiotic cells in anther tissues. To visualize the signals of CUT&Tag, RPGC normalized bigWig files were generated using the bamCoverage option from Deeptools^57^ (v3.5.6). Motif enrichment analysis of TCX5 binding sites was conducted using findMotifsGenome.pl from HOMER, identifying motifs enriched with *p*-value<0.05, which was determined by binomial test.

### Image processing and HEI10 and RAD51 foci quantification

For image analysis of RAD51 and HEI10 immunostainings, maximum intensity projections were generated using the *Z Project* function in Fiji/ImageJ (Version 2.14.0/1.54f). RAD51and HEI10 foci were manually counted on merged channels. Only RAD51 and HEI10 foci colocalizing with the chromosome markers (ASY1 and ZYP1, respectively) were included in the analysis. To ensure consistency across samples, all images were adjusted with identical brightness and contrast settings.

### Statistical analysis

All statistical analyses (as indicated in the figure legends) were performed using GraphPad Prism software (GraphPad Software Version 10.4.1). Error bars indicate standard deviations.

## Data Availability

The RNA-seq and CUT&TAG data have been deposited in GEO under GSE302287 and GSE302286, respectively.

## Acknowledgements

We thank Maren Heese for critical reading of the manuscript. This work was supported by a Pro-Exzellenzia Plus fellowship (H.T.) funded by the European Union and the Free and Hanseatic City of Hamburg (FHH), and by the Deutsche Forschungsgemeinschaft (DFG) (SCHN 736/16-1 to A.S.).

## Author Contributions

H.T. conceived the study, designed the experiments, performed investigations, conducted formal analyses, curated data, prepared visualizations, and wrote the original draft. X.M. contributed to investigations, formal analyses, data curation, and visualizations. L.L. and F.B. contributed to the investigation. D.L. contributed to investigation and supervision. M.B. and A.S. contributed to conceptualization, methodology, supervision, and funding acquisition. A.S. also contributed to writing the original draft. All authors read, commented on and approved the manuscript.

## Extended Data Figure & Table legends

**Extended Data Fig. 1:** DREAM mutants exhibit incomplete synapsis and form dyads and triads; however, DSB formation occurs at WT level. **(A)** Synapsis is incomplete in *aly1/2* and *tcx5/6* mutants. Immunostaining of ASY1 (green) and ZYP1 (magenta) at late pachytene in WT, *aly1/2*, and *tcx5/6* reveals that ASY1 is largely removed from chromosomes, except at rDNA regions, and ZYP1 fully decorates the chromosome axes in WT. In contrast, both ASY1 removal and ZYP1 loading are incomplete in *aly1/2* and *tcx5/6*. **(B)** At the end of meiosis, DREAM mutants more frequently produce triads and dyads instead of normal tetrads, compared to WT. **(C, D)** DSB formation is not impaired in *aly1/2* and *tcx5/6*. **(C)** Immunostaining of RAD51 (magenta, left panel) and ASY1 (green, merged with RAD51 at right panel) at zygotene in WT, *aly1/2* and *tcx5/6*. **(D)** Quantification of RAD51 foci shows no significant difference between WT, *aly1/2*, and *tcx5/6*. Statistical analysis was performed using one-way ANOVA followed by Dunnett’s multiple comparison test. Scale bars, 2 µm.

**Extended Data Fig. 2:** ALY1 and TCX5 are present in meiosis and localize to chromosomes. **(A–B)** Confocal images showing the localization patterns of EGFP-tagged ALY1 (green) **(A)**, TCX5 (green) **(B)**, alongside RFP-tagged TUA5 (magenta), which serves as a meiotic stage marker. Scale bars, 10 µm.

**Extended Data Fig. 3:** ALY1-EGFP and EGFP-TCX5 reporter lines rescue fertility defects in the respective double mutants. **(A)** Representative images from chromosome spreads showing meiotic progression in *ALY1-EGFP aly1/2* and *EGFP-TCX5 tcx5/6*. Scale bars, 10 µm. **(B)** Quantification of bivalents at metaphase I in WT, *aly1/2*, *ALY1-EGFP aly1/2*, *tcx5/6*, and *EGFP-TCX5 tcx5/6* indicates that complementation with ALY1 and TCX5 nearly completely restores proper bivalent formation. Frequency of observed bivalence numbers and chromosome entanglement at metaphase I. Above the chart, “n” indicates the number of metaphase I cells analyzed and the average bivalence number per genotype is given.

**Extended Data Fig. 4:** GO enrichment analysis of downregulated genes in *aly1/2* **(A)** and in *tcx5/6* **(B)** male meiocytes compared to WT.

**Extended Data Fig. 5:** Integration of transcriptomic and CUT&Tag data reveals direct TCX5 targets among misregulated genes in *aly1/2* and *tcx5/6*. **(A)** Heatmap representation of high-confidence EGFP-TCX5 CUT&Tag peaks across gene bodies and flanking regions of upregulated (pink, *n*=271), downregulated (blue, *n*=98), and non-differentially expressed (orange, *n*=3101) genes in *tcx5/6* male meiocytes. Also shown are genes classified as TCX5-non-targeted that are differentially expressed in *tcx5/6*, indicating a stringent threshold used for CUT&Tag data evaluation. Enrichment signals cluster near TSS across all categories. **(B)** Venn diagram illustrating the intersection between genes upregulated (pink) or downregulated (blue) in *aly1/2* male meiocytes and TCX5-bound loci identified by CUT&Tag using EGFP-TCX5 in anthers. Statistical significance was assessed using a hypergeometric test (*p*=1.2e−16 and *p*=0.24). **(C)** Venn diagram depicting the overlap of TCX5 targets that are transcriptionally upregulated in both *tcx5/6* (dark pink) and *aly1/2* (light pink). Statistical significance of the overlap was assessed using a hypergeometric test (*p*=1.7e-236). **(D)** GO enrichment analysis of TCX5-bound genes that are consistently upregulated in *tcx5/6* and *aly1/2* male meiocytes.

**Extended Data Fig. 6:** A *MER3-EGFP* genomic reporter under the meiosis-specific *DMC1* promoter produces functional MER3 protein and rescues CO defects in *mer3*. **(A)** Cytological analysis of meiotic progression in *PRO_DMC1_:MER3-EGFP mer3* and *mer3* single mutant via chromosome spreads. **(B)** Confocal images of MER3-EGFP at zygotene, pachytene, after nuclear envelope breakdown (NEB), and telophase II in *mer3*, *aly1/2*, and *tcx5/6*. First row: MER3–EGFP (green). Second row: merged GFP with brightfield (gray) and autofluorescence (dark blue). MER3-EGFP is expressed specifically in meiotic cells within anthers in *mer3* but shows ectopic expression in tapetal cells of *aly1/2* and *tcx5/6*. The localization within meiotic cells is comparable across genotypes. Scale bars, 10 µm.

**Extended Data Fig. 7:** Comparative analysis of TCX5 targets in somatic and reproductive tissues. **(A)** Venn diagram of TCX5-bound genes identified by ChIP–seq in seedlings^4^ (light green) and by CUT&Tag in anthers (light yellow), highlighting shared and tissue-specific binding (*p*=9.0e-315). **(B)** Venn diagram showing the overlap between TCX5 targets identified in anthers that are upregulated in *tcx5/6* meiocytes and TCX5 targets identified in seedlings that are upregulated in *tcx5/6* seedlings^4^ (*p*=3.1e-150). **(C)** GO enrichment of genes bound by TCX5 in both tissues and upregulated in *tcx5/6* meiocytes and seedlings**. (D)** Venn diagram of upregulated (pink) and downregulated (blue) genes in *tcx5/6* seedlings relative to TCX5-bound loci identified in seedlings (*p*=5.0e−228 and *p*=0.79, respectively). **(E)** Overlap between TCX5 targets identified in anthers that are downregulated in *tcx5/6* meiocytes and TCX5 targets identified in seedlings that are downregulated in *tcx5/6* seedlings (*p*=3.1e-150). Statistical significance in A, B, and D was assessed by a hypergeometric test. Seedling transcriptome and TCX5–FLAG ChIP–seq data are from Ning *et al.* (2020) and were reanalyzed here.

**Extended Data Table 1:** Transmission efficiency analysis of the *tcx6* allele in a *tcx5* mutant background indicates no gametophytic defects. Reciprocal crosses were performed between WT and plants homozygous for *tcx5* and heterozygous for *tcx6*. F□ progeny were genotyped to assess segregation of the *TCX6* mutant and WT alleles. A transmission efficiency of 100% indicates equal transmission of the two alleles (i.e., 50% of the FL seedlings are heterozygous). Values greater than 100% for one allele necessarily reflect reduced transmission of the other, and vice versa. Results show that the *tcx5 tcx6* mutation combination does not impair gamete transmission.

**Extended Data Table 2:** Sequencing quality metrics for RNA-seq of *aly1/2*, *tcx5/6*, and WT male meiocytes. Summary of whole transcriptome sequencing statistics, including the number of raw reads, total base counts, clean reads, error rate, GC content, and the proportion of bases falling within Q20 and Q30 quality thresholds, corresponding to base call accuracies of 99% and 99.9%, respectively.

**Extended Data Table 3:** List of 135 meiotic genes and their transcriptional regulation by the DREAM complex in somatic and reproductive tissues. Differential expression of meiotic genes in *aly1/2* and *tcx5/6* male meiocytes compared to WT is shown as log□ fold change values with adjusted *P* values. The table also indicates TCX5 target status (–, not targeted; +, TCX5 target if TCX5 enrichment fold-change>2 in both replicates) based on CUT&Tag in anthers (this study) and ChIP– seq in seedlings (Ning *et al.,* 2020), as well as their expression in *tcx5/6* seedlings (Ning *et al.*, 2020). Expression changes are color-coded: gray, |log□FoldChange|<0.5; black, not significant (*p*adj≥0.05); brown, significantly upregulated (log□FoldChange≥0.5, *p*adj<0.05); blue, significantly downregulated (log□FoldChange≤–0.5, *p*adj<0.05). The final column assigns each gene to one of the regulatory classes defined in Fig. 4A, based on their transcriptional regulation in *tcx5/6* meiocytes and TCX5 binding status in anthers and seedlings.

## References

1. Fischer, M. & Müller, G. A. Cell cycle transcription control: DREAM/MuvB and RB-E2F complexes. Crit. Rev. Biochem. Mol. Biol. 52, 638–662 (2017).

2. Fischer, M., Schade, A. E., Branigan, T. B., Müller, G. A. & DeCaprio, J. A. Coordinating gene expression during the cell cycle. Trends Biochem. Sci. 47, 1009–1022 (2022).

3. Lang, L. et al. The DREAM complex represses growth in response to DNA damage in Arabidopsis. Life Sci. Alliance 4, e202101141 (2021).

4. Ning, Y. Q., et al. DREAM complex suppresses DNA methylation maintenance genes and precludes DNA hypermethylation. Nat. Plants 4, (2020).

5. Koliopoulos, M. G. et al. Structure of a nucleosome-bound MuvB transcription factor complex reveals DNA remodelling. Nat. Commun. 13, 5075 (2022).

6. Sadasivam, S., Duan, S. & DeCaprio, J. A. The MuvB complex sequentially recruits B-Myb and FoxM1 to promote mitotic gene expression. Genes Dev. 26, 474–489 (2012).

7. Fischer, M., Grossmann, P., Padi, M. & DeCaprio, J. A. Integration of TP53, DREAM, MMB-FOXM1 and RB-E2F target gene analyses identifies cell cycle gene regulatory networks. Nucleic Acids Res. 44, 6070–6086 (2016).

8. Kobayashi, K. et al. Transcriptional repression by MYB3R proteins regulates plant organ growth. EMBO J. 34, 1992–2007 (2015).

9. Wang, Y. et al. The Arabidopsis DREAM complex antagonizes WDR5A to modulate histone H3K4me2 / 3 deposition for a subset of genome repression. Proc. Natl. Acad. Sci. 1–9 (2022)

10. Lang, L. et al. The two plant-specific DREAM components FLIC and FLAC repress floral transition in Arabidopsis. 2023.08.07.552284 Preprint at 10.1101/2023.08.07.552284 (2023).

11. Bouyer, D., et al. Genome-Wide Identification of RETINOBLASTOMA RELATED 1 Binding Sites in Arabidopsis Reveals Novel DNA Damage Regulators. PLoS Genetics vol. 14 (2018).

12. Keeney, S., Giroux, C. N. & Kleckner, N. Meiosis-Specific DNA Double-Strand Breaks Are Catalyzed by Spo11, a Member of a Widely Conserved Protein Family. Cell 88, 375–384 (1997).

13. Grelon, M., Vezon, D., Gendrot, G. & Pelletier, G. AtSPO11L1 is necessary for efficient meiotic recombination in plants. EMBO J. 20, 589–600 (2001).

14. Armstrong, S. J., Caryl, A. P., Jones, G. H. & Franklin, F. C. H. Asy1, a protein required for meiotic chromosome synapsis, localizes to axis-associated chromatin in Arabidopsis and Brassica. J. Cell Sci. 115, 3645–3655 (2002).

15. Ferdous, M. et al. Inter-Homolog Crossing-Over and Synapsis in Arabidopsis Meiosis Are Dependent on the Chromosome Axis Protein AtASY3. PLoS Genet. 8, e1002507 (2012).

16. Ines, O. D. et al. Meiotic Recombination in Arabidopsis Is Catalysed by DMC1, with RAD51 Playing a Supporting Role. PLOS Genet. 9, e1003787 (2013).

17. Cloud, V., Chan, Y.-L., Grubb, J., Budke, B. & Bishop, D. K. Rad51 is an accessory factor for Dmc1-mediated joint molecule formation during meiosis. Science 337, 1222–1225 (2012).

18. Zickler, D. & Kleckner, N. Recombination, Pairing, and Synapsis of Homologs during Meiosis. Cold Spring Harb. Lab. Press 1,2 (2015).

19. Baudat, F. & de Massy, B. Regulating double-stranded DNA break repair towards crossover or non-crossover during mammalian meiosis. Chromosome Res. 15, 565–577 (2007).

20. Bishop, D. K. & Zickler, D. Early Decision: Meiotic Crossover Interference prior to Stable Strand Exchange and Synapsis. Cell 117, 9–15 (2004).

21. Chen, C., Zhang, W., Timofejeva, L., Gerardin, Y. & Ma, H. The Arabidopsis ROCK-N-ROLLERS gene encodes a homolog of the yeast ATP-dependent DNA helicase MER3 and is required for normal meiotic crossover formation. Plant J. 43, 321–334 (2005).

22. Lu, P., Wijeratne, A. J., Wang, Z., Copenhaver, G. P. & Ma, H. Arabidopsis PTD Is Required for Type I Crossover Formation and Affects Recombination Frequency in Two Different Chromosomal Regions. J. Genet. Genomics 41, 165–175 (2014).

23. Ren, Y. et al. OsSHOC1 and OsPTD1 are essential for crossover formation during rice meiosis. Plant J. 98, 315–328 (2019).

24. Chelysheva, L. et al. Zip4/Spo22 is required for class I CO formation but not for synapsis completion in Arabidopsis thaliana. PLoS Genet. 3, 802–813 (2007).

25. Higgins, J. D. et al. AtMSH5 partners AtMSH4 in the class I meiotic crossover pathway in Arabidopsis thaliana, but is not required for synapsis. Plant J. 55, 28– 39 (2008).

26. Chelysheva, L. et al. The Arabidopsis HEI10 is a new ZMM protein related to Zip3. PLoS Genet. 8, (2012).

27. Sadasivam, S. & DeCaprio, J. A. The DREAM complex: Master coordinator of cell cycle-dependent gene expression. Nat. Rev. Cancer 13, 585–595 (2013).

28. Girard, C. et al. FANCM-associated proteins MHF1 and MHF2, but not the other Fanconi anemia factors, limit meiotic crossovers. Nucleic Acids Res. 42, 9087– 9095 (2014).

29. Higgins, J. D., Ferdous, M., Osman, K. & Franklin, F. C. H. The RecQ helicase AtRECQ4A is required to remove inter-chromosomal telomeric connections that arise during meiotic recombination in Arabidopsis. Plant J. 65, 492–502 (2011).

30. Séguéla-Arnaud, M. et al. Multiple mechanisms limit meiotic crossovers: TOP3α and two BLM homologs antagonize crossovers in parallel to FANCM. Proc. Natl. Acad. Sci. 112, 4713–4718 (2015).

31. Crismani, W. et al. FANCM Limits Meiotic Crossovers. Science 336, 1588–1590 (2012).

32. Serra, H. et al. Massive crossover elevation via combination of HEI10 and recq4a recq4b during Arabidopsis meiosis. 159764 Preprint at 10.1101/159764 (2017).

33. Mercier, R. et al. Two Meiotic Crossover Classes Cohabit in Arabidopsis: One Is Dependent on MER3,whereas the Other One Is Not. Curr. Biol. 15, 692–701 (2005).

34. Georlette, D. et al. Genomic profiling and expression studies reveal both positive and negative activities for the Drosophila Myb–MuvB/dREAM complex in proliferating cells. Genes Dev. 21, 2880–2896 (2007).

35. Goetsch, P. D., Garrigues, J. M. & Strome, S. Loss of the Caenorhabditis elegans pocket protein LIN-35 reveals MuvB’s innate function as the repressor of DREAM target genes. PLOS Genet. 13, e1007088 (2017).

36. Tabuchi, T. M. et al. Chromosome-biased binding and gene regulation by the caenorhabditis elegans DRM complex. PLoS Genet. 7, 1002074 (2011).

37. Bujarrabal-Dueso, A. et al. The DREAM complex functions as conserved master regulator of somatic DNA-repair capacities. Nat. Struct. Mol. Biol. 30, 475–488 (2023).

38. Bleuyard, J.-Y. & White, C. I. The Arabidopsis homologue of Xrcc3 plays an essential role in meiosis. EMBO J. 23, 439–449 (2004).

39. Grelon, M., Gendrot, G., Vezon, D. & Pelletier, G. The Arabidopsis MEI1 gene encodes a protein with five BRCT domains that is involved in meiosis-specific DNA repair events independent of SPO11-induced DSBs. Plant J. Cell Mol. Biol. 35, 465–475 (2003).

40. d’Erfurth, I. et al. Mutations in AtPS1 (Arabidopsis thaliana Parallel Spindle 1) Lead to the Production of Diploid Pollen Grains. PLoS Genet. 4, e1000274 (2008).

41. Laktionov, P. P. et al. Genome-wide analysis of gene regulation mechanisms during Drosophila spermatogenesis. Epigenetics Chromatin 11, 1–15 (2018).

42. Tabuchi, T. M., Rechtsteiner, A., Strome, S. & Hagstrom, K. A. Opposing Activities of DRM and MES-4 Tune Gene Expression and X-Chromosome Repression in Caenorhabditis elegans Germ Cells. G3 GenesGenomesGenetics 4, 143–153 (2013).

43. Jiang, J., Benson, E., Bausek, N., Doggett, K. & White-Cooper, H. Tombola, a tesmin/TSO1-family protein, regulates transciptional activation in the Drosophila male germline and physically interacts with always early. Development 134, 1549–1559 (2007).

44. Oji, A. et al. TESMIN, METALLOTHIONEIN-LIKE 5, is required for spermatogenesis in mice†. Biol. Reprod. 1–27 (2020) doi:10.1093/biolre/ioaa002.

45. Zhang, X. et al. Nuclear translocation of MTL5 from cytoplasm requires its direct interaction with LIN9 and is essential for male meiosis and fertility. PLOS Genet. 837 17, e1009753 (2021).

46. Komaki, S. & Schnittger, A. The Spindle Assembly Checkpoint in Arabidopsis Is Rapidly Shut Off during Severe Stress. Dev. Cell 43, 172-185.e5 (2017).

47. Yang, C. et al. The Arabidopsis Cdk1/Cdk2 homolog CDKAL;1 controls chromosome axis assembly during plant meiosis. EMBO J. 1–19 (2019) doi:10.15252/embj.2019101625.

48. Nakagawa, T. et al. Improved gateway binary vectors: High-performance vectors for creation of fusion constructs in transgenic analysis of plants. Biosci. Biotechnol. Biochem. 71, 2095–2100 (2007).

49. Zhang, X., Henriques, R., Lin, S.-S., Niu, Q.-W. & Chua, N.-H. Agrobacterium-mediated transformation of Arabidopsis thaliana using the floral dip method. Nat. Protoc. 1, 641–646 (2006).

50. Peterson, R., Slovin, J. P. & Chen, C. A Simplified Method for Differential Staining of Aborted and Non-Aborted Pollen Grains. *Int*. J. Plant Biol. 1, e13 (2010).

51. Sims, J., Schlögelhofer, P. & Kurzbauer, M. T. From Microscopy to Nanoscopy: Defining an Arabidopsis thaliana Meiotic Atlas at the Nanometer Scale. Front. Plant Sci. 12, (2021).

52. Kurzbauer, M.-T., Uanschou, C., Chen, D. & Schlögelhofer, P. The recombinases DMC1 and RAD51 are functionally and spatially separated during meiosis in Arabidopsis. Plant Cell 24, 2058–2070 (2012).

53. Tian, T. et al. agriGO v2.0: a GO analysis toolkit for the agricultural community, 2017 update. Nucleic Acids Res. 45, W122–W129 (2017).

54. Langmead, B. & Salzberg, S. L. Fast gapped-read alignment with Bowtie 2. Nat. Methods 9, 357–359 (2012).

55. Li, H. et al. The Sequence Alignment/Map format and SAMtools. Bioinformatics 25, 2078–2079 (2009).

56. Zhang, Y. et al. Model-based Analysis of ChIP-Seq (MACS). Genome Biol. 9, R137 (2008).

57. Ramírez, F., Dündar, F., Diehl, S., Grüning, B. A. & Manke, T. deepTools: a flexible platform for exploring deep-sequencing data. Nucleic Acids Res. 42, W187–W191 (2014).

