## Supplementary figures and images for "Transcriptional control of meiotic recombination by the DREAM complex"

### Supplemental figure 1

**Extended Data Fig. 1**

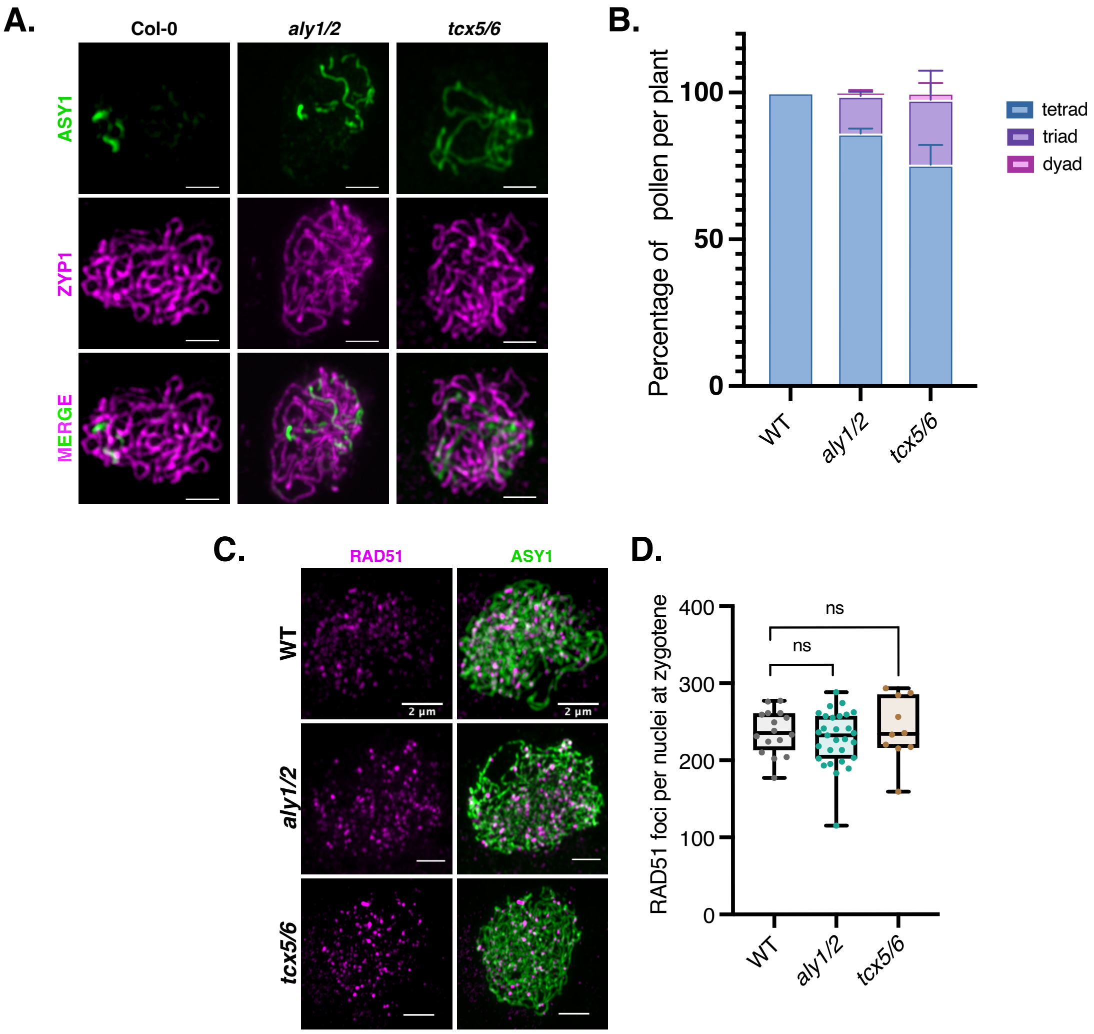

### Supplemental figure 2

#### Extended Data Fig. 2

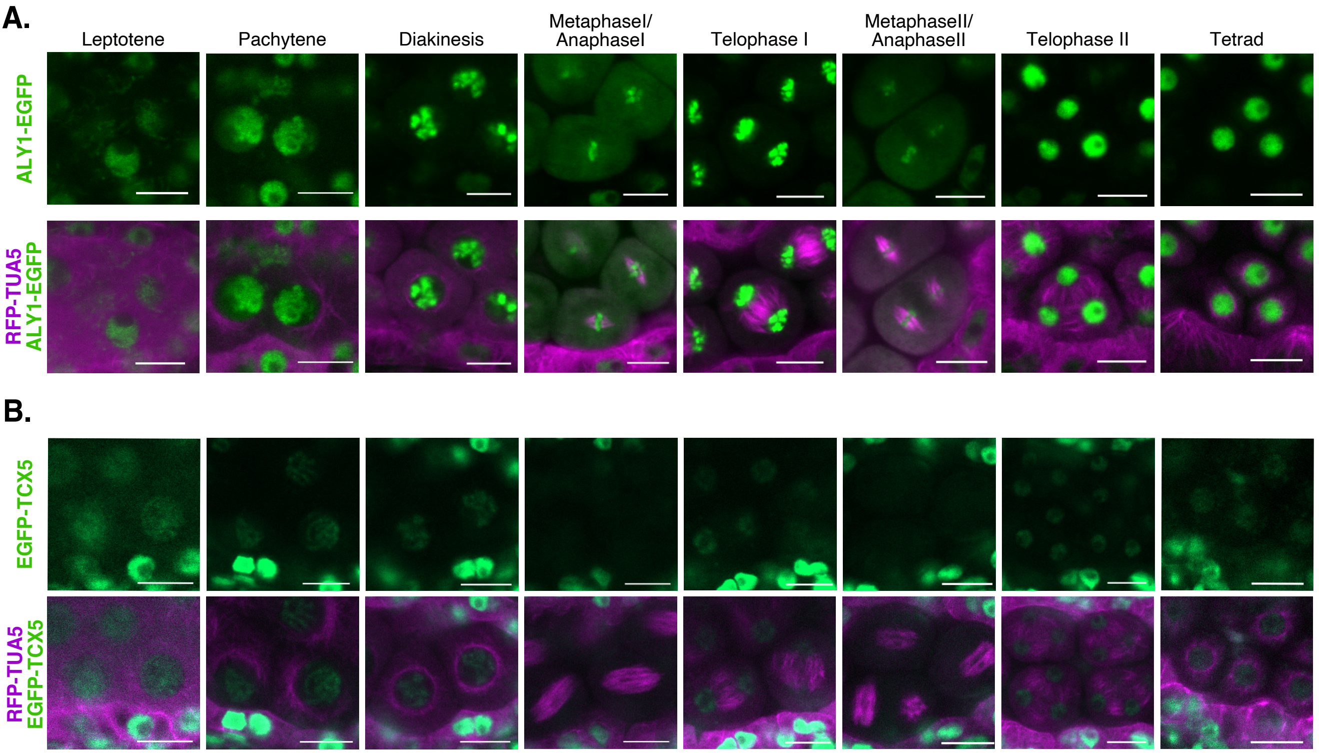

### Supplemental figure 3

#### Extended Data Fig. 3

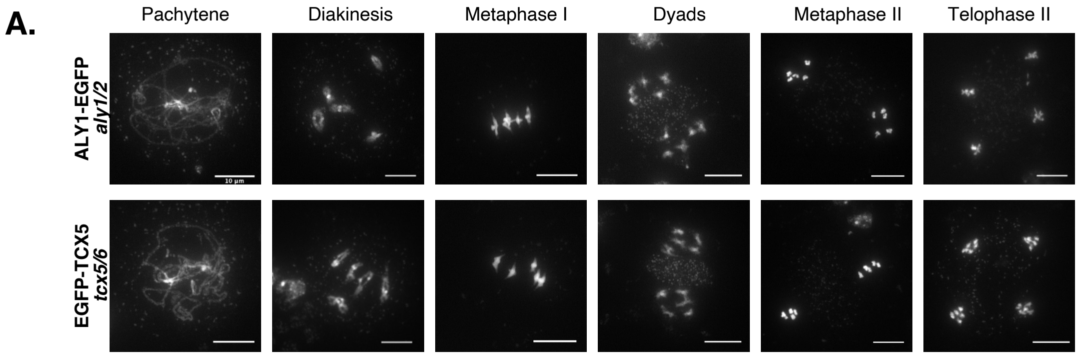


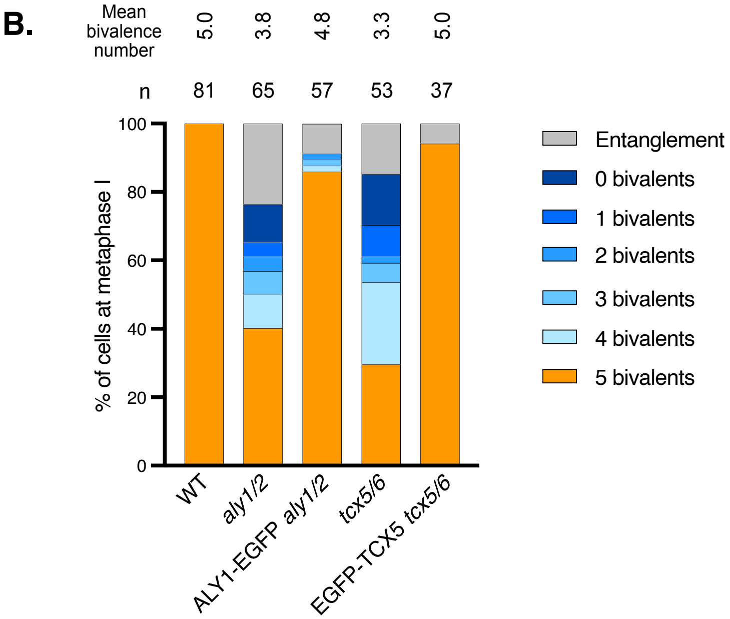

### Supplemental figure 4

##### Extended Data Fig. 4

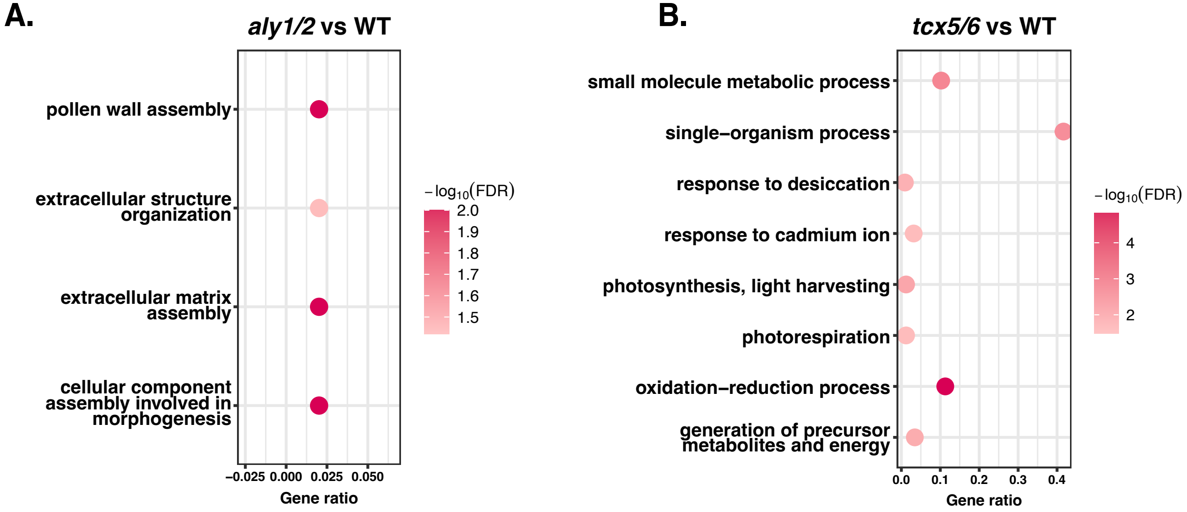

### Supplemental figure 5

#### Extended Data Fig. 5

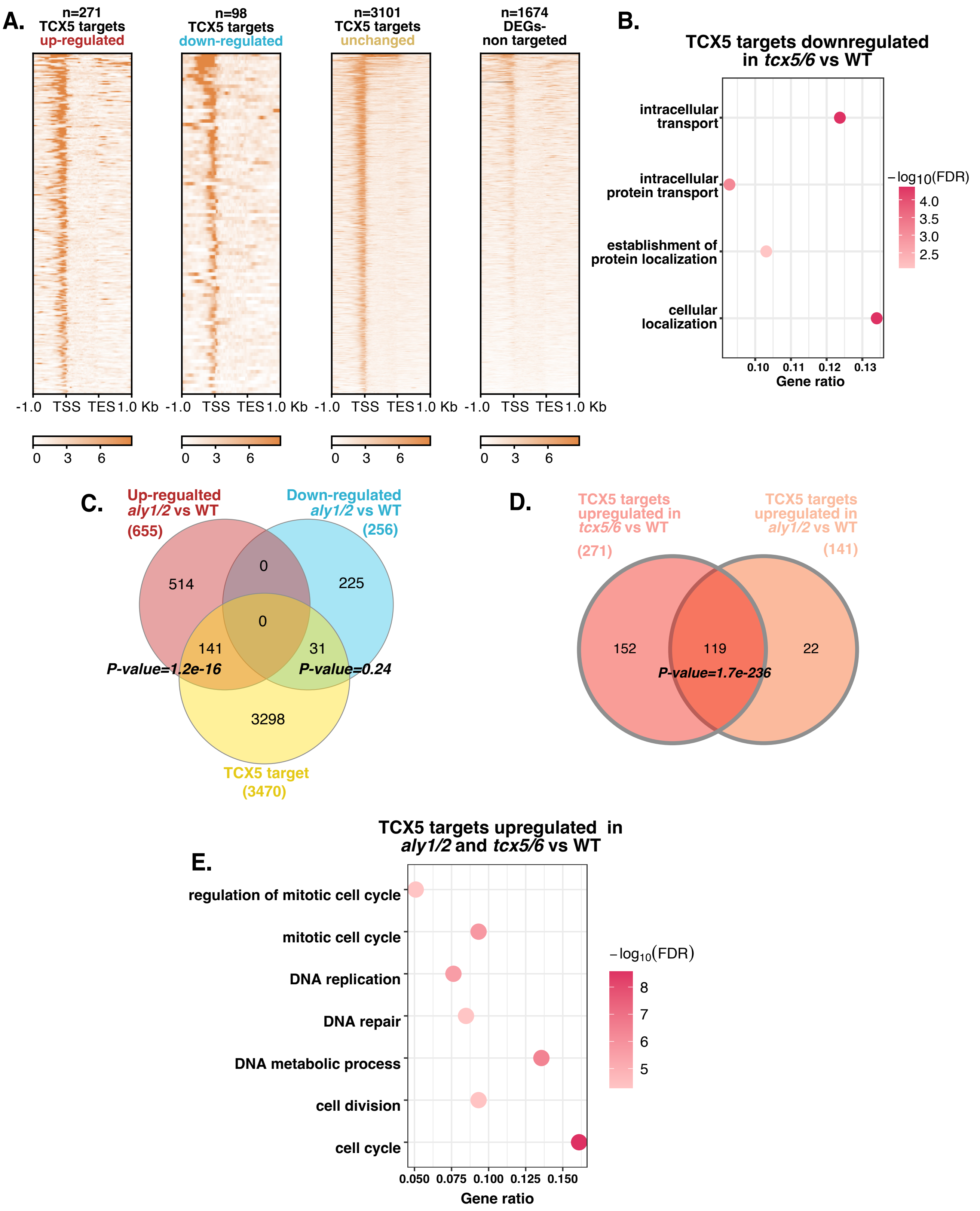

### Supplemental figure 6

##### Extended Data Fig. 6

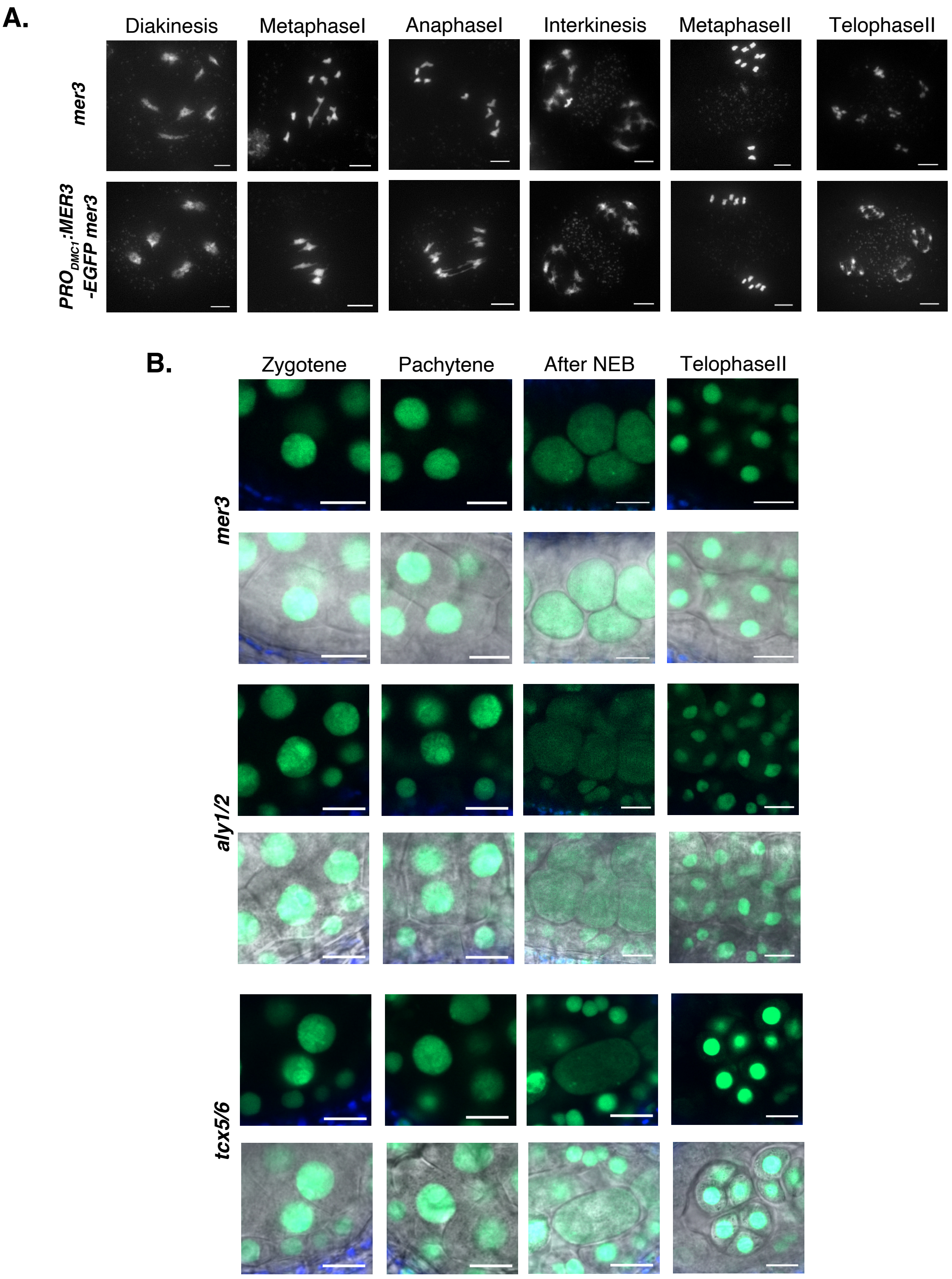

### Supplemental figure 7

##### Extended Data Fig. 7

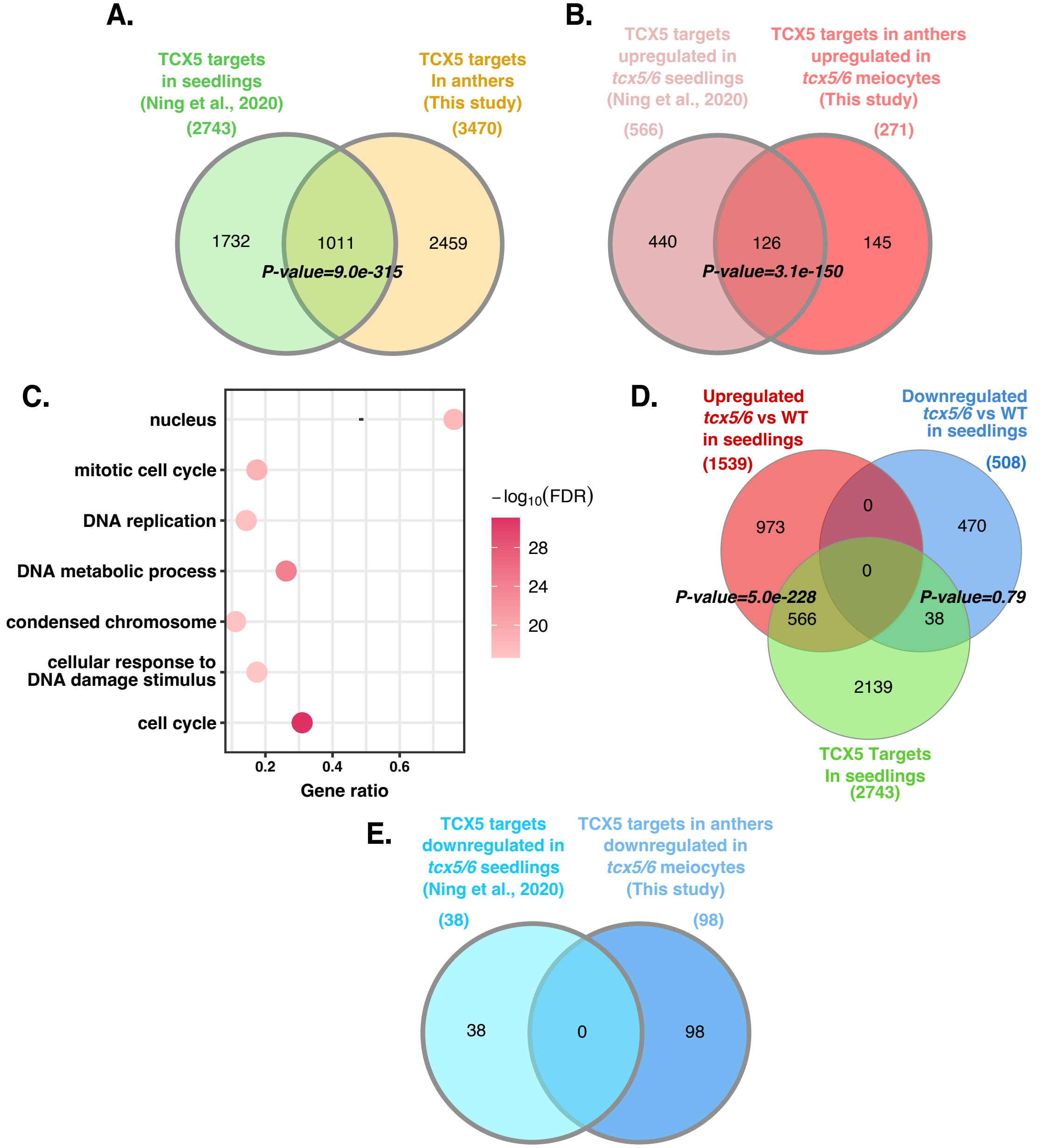
